# Generating genomic platforms to study *Candida albicans* pathogenesis

**DOI:** 10.1101/261628

**Authors:** Mélanie Legrand, Sophie Bachellier-Bassi, Keunsook K. Lee, Yogesh Chaudhari, Hélène Tournu, Laurence Arbogast, Hélène Boyer, Murielle Chauvel, Vitor Cabral, Corinne Maufrais, Audrey Nesseir, Irena Maslanka, Emmanuelle Permal, Tristan Rossignol, Louise A. Walker, Ute Zeidler, Sadri Znaidi, Floris Schoeters, Charlotte Majgier, Renaud A. Julien, Laurence Ma, Magali Tichit, Christiane Bouchier, Patrick Van Dijck, Carol A. Munro, Christophe d’Enfert

**Author notes:** The authors wish it to be known that, in their opinion, the first two authors should be regarded as joint First Authors. Corresponding authors: Christophe d’Enfert and Carol Munro, To whom correspondence should be addressed. Correspondence may also be addressed to Carol Munro. Present Address: Yogesh Chaudhari, Biosciences Department, University of Exeter, Exeter, EX4 4QD, UK; Hélène Tournu, Department of Clinical Pharmacy, University of Tennessee, Memphis, TN, 38163, USA; Laurence Arbogast, Brain plasticity in response to the environment Group, Institut Pasteur, Paris, 75015, France; Hélène Boyer, Laboratoire d’Immunologie, Hôpital Henri Mondor, Créteil, 94000, France; Vitor Cabral, Bacterial Signalling Group, Instituto Gulbenkian de Ciência, Oeiras, 2780-156, Portugal; Audrey Nesseir, Lycée Grégor Mendel, Vincennes, 94300, France; Emmanuelle Permal, Toulouse, 31000, France; Tristan Rossignol, MICALIS, INRA, Jouy-enJosas, 78352, France; Ute Zeidler, Sandoz, Unterach, 4866, Austria; Sadri Znaidi, Laboratory of Molecular Microbiology, Vaccinology & Biotechnology Development, Institut Pasteur de Tunis, Tunis-Belvédère, 1002, Tunisia; Magali Tichit, Laboratoire Histopathologie Humaine et Modèles Animaux, Institut Pasteur, Paris, 75015, France.

## Abstract

The advent of the genomic era has made elucidating gene function at large scale a pressing challenge. ORFeome collections, whereby almost all ORFs of a given species are cloned and can be subsequently leveraged in multiple functional genomic approaches, represent valuable resources towards this endeavor. Here we provide novel, genome-scale tools for the study of *Candida albicans*, a commensal yeast that is also responsible for frequent superficial and disseminated infections in humans. We have generated an ORFeome collection composed of 5,102 ORFs cloned in a Gateway^™^ donor vector, representing 83% of the currently annotated coding sequences of *C. albicans*. Sequencing data of the cloned ORFs are available in the CandidaOrfDB database at http://candidaorfeome.eu. We also engineered 49 expression vectors with a choice of promoters, tags, and selection markers and demonstrated their applicability to the study of target ORFs transferred from the *C. albicans* ORFeome. In addition, the use of the ORFeome in the detection of protein-protein interaction was demonstrated. Mating-compatible strains as well as Gateway^™^-compatible two-hybrid vectors were engineered, validated and used in a proof of concept experiment. These unique and valuable resources should greatly facilitate future functional studies in *C. albicans* and the elucidation of mechanisms that underlie its pathogenicity.

## INTRODUCTION

Over the last decade, there has been an exponential growth in the quantity of available genome sequence data due to the very rapid progress in sequencing technology. In 2004, the genome sequence of the human fungal pathogen *Candida albicans* was released as Assembly 19 (1). With the challenge of working with a heterozygous diploid organism, new computational methods had to be developed and resulted in the release in 2013 of Assembly 22, an assembly of a completely phased diploid genome sequence for the standard *C. albicans* reference strain SC5314 (2). This opened new perspectives to understand the genetic basis and functional mechanisms that underlie pathogenesis and evolution in this organism. Although *C. albicans* gene sequences have been available to the community for more than a decade (3,4), only 1,670 out of the 6,198 predicted protein-coding genes have been characterized as of January 31st, 2018 according to the Candida Genome Database (5). Nowadays, the growing availability of whole-genome datasets has encouraged a shift towards the development of functional genomics and systems biology, enabling analysis of high-throughput whole-genome assays to better understand biological networks. In this context, major efforts have been made to generate large-scale deletion (6-9) or overexpression (10-12) mutant collections. Genome-wide ORF libraries or ORFeomes represent useful resources for the implementation of approaches used to elucidate gene function (13). Besides the development of collections of overexpression mutants (11), ORFeomes facilitate approaches aimed at evaluating protein subcellular localization and identifying protein-protein interactions (yeast two-hybrid) both at steady state and in response to environmental stimuli (14).

Large-scale cloning projects, with the goal of cloning all predicted ORFs into flexible recombinational vectors, have been described for several model organisms, including *Caenorhabditis elegans, Saccharomyces cerevisiae, Schizosaccharomyces pombe*, *Escherichia coli* K-12, *Arabidopsis thaliana*, *Xenopus laevis* and *Drosophila melanogaster*, as well as infectious microorganisms such as *Brucella melitensis, Plasmodium falciparum*, *Helicobacter pylori*, *Chlamydia pneumonia*, *Staphylococcus aureus* and viruses (15-28). Human ORFeomes have also been generated and made publicly available, with the latest version containing sequence-confirmed ORFs for more than 11,000 human genes (29). Most ORFeome libraries are created using the highly versatile Gateway^™^ cloning approach (30). The Gateway^™^ technology employs the recombination system of bacteriophage lambda with two sets of reactions: the BP reaction, catalysed by Gateway^™^ BP Clonase^™^, facilitates recombination between *attB* and *attP* sequences, and the LR reaction, catalysed by Gateway^™^ LR Clonase^™^, promotes recombination between *attL* and *attR* sequences (30). Previously, we have reported the development of a first generation *C. albicans* ORFeome, in which 644 full-length ORFs, encoding signaling proteins (transcription factors and kinases), as well as cell wall proteins and proteins involved in DNA processes (repair, replication and recombination), were cloned in the Gateway^™^ vector pDONR207 (31). Here we describe the generation and validation of the latest *Candida albicans* ORFeome collection, which consists of 5,102 ORFs in pDONR207, allowing the transfer of the cloned genes into a variety of *C. albicans* Gateway^™^-compatible expression vectors. To this end, we also provide a panel of 15 expression vectors that differ in the combination of promoter/selection marker they are carrying, and give different options regarding the tagging of the cloned gene product. These 15 expression vectors have been validated using the filament-specific transcription factor *UME6*. In addition, we also provide a secondary collection of 34 expression vectors with additional options regarding the choice of the promoter and the tagging of the cloned gene product. Finally, among the numerous applications of ORFeome collections, we present a powerful association of the *C. albicans* ORFeome and a *C. albicans*-adapted two-hybrid system (32) that will allow systematic protein-protein interaction screening in *C. albicans*.

## MATERIAL AND METHODS

### Strains and growth conditions

*C. albicans* strains used in this study are listed in **Table 1**. *C. albicans* strains were routinely cultured at 30°C in YPD medium (1% yeast extract, 2% peptone, 2% dextrose), synthetic dextrose (SD) medium (0.67% yeast nitrogen base, 2% dextrose) or nourseothricin-containing YPD medium (YPD + 200 μg/ml Nourseothricin). Solid media were obtained by adding 2% agar.

*E. coli* strains were routinely cultured at 30°C or 37°C in LB or 2YT supplemented with 10 μg/ml gentamicin, 50 μg/ml kanamycin or 50 μg/ml ticarcillin. Solid media were obtained by adding 2% agar.

DH5α *E. coli* cells competent for transformation with DNA were prepared according to Hanahan *et al*. (33).

The ORFeome collection is being distributed by the Centre International de Ressources Microbiennes (CIRM). The 49 expression vectors are available from Addgene, while two-hybrid strains and vectors are now accessible from the Belgian Co-ordinated Collections of Micro-organisms (BCCM).

### Generation of an entry clone collection

#### Primer design

The DNA sequence of *C. albicans* was obtained from the Candida Genome Database Release 21 (www.candidagenome.org), before Assembly 22 was released. Gene-specific forward primers were designed by adding the sequence 5’-GGGGACAAGTTTGTACAAAAAAGCAGGCTTG-3’ to the 5’ end of the first 30 nucleotides of each ORF. Gene-specific reverse primers were designed by adding the sequence 5’-GGGGACCACTTTGTACAAGAAAGCTGGGTC-3’ to the 5’ end of the last 30 nucleotides of each ORF excluding the Stop codon. PCR fragments amplified from these primers are compatible for C-terminal tagging with destination vectors containing the Gateway^™^ cassette A. A total of 6,205 primer pairs were obtained from Invitrogen in a 96-well format and are listed in **Supplemental Table S1**.

#### Gateway^™^ cloning of the C. albicans ORFeome

The detailed method for the cloning of *C. albicans* ORFs in the pDONR207 vector has been described (34). Briefly, ORFs, ranging from 90 bp to 5,295 bp, were amplified from genomic DNA of *C. albicans* strain SC5314 in 96-well plates using the Thermo Scientific Phusion High-Fidelity DNA Polymerase and 30 cycles of amplification, with elongation time varying from 1 to 3 min according to the ORF size. After ethanol precipitation, Gateway^™^-compatible amplified ORFs were recombined into pDONR207 (Invitrogen) using the Gateway^™^ BP Clonase^™^ II Enzyme Mix (Invitrogen). Reaction mixes containing pDONR207, the PCR products and the BP Clonase^™^ were incubated overnight in 96-well plates, at room temperature. After adding proteinase K (Invitrogen) and incubating 10 min at 37°C, the BP reactions were directly used for bacterial transformation. 45-50 μl of chemically competent *E. coli* DH5α were added to the BP reactions and incubated for 30 min at 4°C. After heat-shock at 42°C for 35 s, 150 μl of SOC medium (20 ml YPD + 2 ml LB + 1 ml Hepes) was added to the transformation reactions and the samples were covered with breathing films and incubated for 1.5 h at 37°C with shaking. Then 100 μl of each sample were plated onto LB agar containing 10 μg/ml Gentamicin and incubated overnight at 37°C. The remainder of the transformation reactions were stored at −80°C in 30% glycerol. A single colony from each transformation reaction was inoculated in 400 μl 2YT + 10 μg/ml Gentamicin in 96 deep-well microplates. After 36h-growth at 37°C, the cultures were used for plasmid extraction and for −80°C storage.

#### Validation of entry clones by DNA sequencing and bioinformatics analysis

Sanger sequencing of the 5’-end of each BP clone was performed with the 207VER-F oligonucleotide (see **Supplemental Table 2** for oligonucleotide sequences except those used to amplify ORFs) to confirm that the expected ORF had been cloned. The 3’-ends were also sequenced with the 207VER-R oligonucleotide to ensure that the oligonucleotides did not carry deletions. All inserts validated by Sanger sequencing were systematically subjected to full-length sequence analysis using Illumina technology as described in Chauvel *et al*. (11). If nonsense or frameshift mutations were observed in a cloned ORF, another colony was checked, or the ORF reamplified and cloned again. If these attempts were unsuccessful, cloning of the ORF was abandoned. Clones containing contiguous deletions or insertions of multiples of 3 bp were accepted. Missense mutations and those located within introns were accepted. It should be noted that further analysis of each mutation-containing ORF is required to determine whether or not mutations affect the function of the ORF.

### CandidaOrfDB database

CandidaOrfDB was created to integrate the data of the *C. albicans* ORFeome project and is available at http://candidaorfeome.eu. The database is a relational database using the SQL database server Oracle with a web interface developed using J2EE. The alignment algorithm uses *biojava3-alignment* Java library, implemented with a NeedlemanWunsch aligner, configured with a gap open penalty of 5 and a gap extension penalty of 2. The substitution matrix used is named “nuc-4_4” and was created by Todd Lowe (https://github.com/sbliven/biojava/blob/master/biojava3-alignment/src/main/resources/nuc-4_4.txt). The alignment algorithm was specifically customized for CandidaOrfDB to work on segments of 500 codons, which provides an optimized alignment performance for *C. albicans* average sequence length.

### Clone access

The *C. albicans* ORFeome clones are available at the Centre International de Ressources Microbiennes (CIRM - https://www6.inra.fr/cirm_eng/).

### Destination plasmids

All destination plasmids constructed in this study were derived from CIp-P*_TET_*-GTW (11) and were confirmed by Sanger sequencing. Plasmid sequences have been submitted to GenBank. Accession numbers are given in **Table 2**. All oligonucleotides used for PCR and/or sequencing are listed in **Supplemental Table 2**.

#### SP cloning

First, a spacer sequence (SP) was amplified from the *E. coli kanR* gene borne on pCR-topo-blunt plasmid (Invitrogen) with primers SP3 and SP4. The PCR product was cloned in the SacII site of CIp-P*_TET_*-GTW, yielding CIp-P*_TET_*-GTW-SP, also referred to as pCA-Dest1100. This 390 bp-long sequence is flanked by the SP1 and SP2 Illumina paired-end sequencing primers 1 and 2 and can be used to insert molecular barcodes if expression plasmids are to be used in signature-tagged mutagenesis approaches.

#### Epitope tags

We then inserted in frame tags either upstream of attR1 or downstream of attR2 to allow N-terminal or C-terminal protein tagging, respectively. For the cloning of N-terminal tags, we used pUC-attR1-CmR, containing a PciI-BglII fragment from CIp-GTW cloned into the PciI and BamHI sites of pUC18. CIp-GTW was obtained after deleting P*_TET_* from CIp-P*_TET_*-GTW with Acc65I and HpaI. For cloning of the 3xHA coding sequence, two oligonucleotides (oligo1_PciHABsrG and oligo2_PciHABsrG) containing three HA epitopes were hybridized, gel purified, and cloned into pUC-attR1-CmR cut with PciI and BsrGI. The resulting plasmid was then used as a PCR template with primers GTW02 and GTW03. The resulting PCR product was cut with BspEI and HpaI, and cloned into the same sites of pCA-Dest1100 to yield pCA-Dest1110. For cloning of the GFP tag, the GFP gene was amplified from pFA-GFP-URA3 (35) with primers GTW04 and GTW05. The PCR product was cut with PciI and EcoRV and ligated into the same sites of pUC-attR1-CmR. The resulting plasmid was then used as a PCR template with primers GTW07 and GTW03. The resulting PCR product was cut with ScaI and BspEI and ligated into pCA-Dest1100 cut with HpaI and BspEI, yielding pCA-Dest1120. For cloning of the TAP-tag, the TAP-tag coding region was PCR amplified from pFA-TAP-URA3 (11) with primers GTW13 and GTW14. The resulting PCR product was then cut with PciI and BsrGI, and ligated into the same sites of pUC-attR1-CmR. The resulting plasmid was then digested with EcoRV and BspEI and the smallest restriction fragment was ligated into pCA-Dest1100 cut with HpaI and BspEI to yield pCA-Dest1130. For the cloning of C-terminal tags, we generated pUC-attR2 by inserting a SalI-HindIII fragment from pCA-Dest1100 into the same sites of pUC18. For cloning of the 3xHA epitope, pCaMPY-3xHA (36) was used as a PCR template with primers GTW20 and GTW21, and the resulting PCR product cut with BsrGI and NsiI was ligated in the same sites of pUC-attR2. The resulting plasmid was cut with SalI and NsiI and the fragment cloned in the same sites of pCA-Dest1100, yielding pCA-Dest1101. For cloning the GFP tag, the GFP gene was amplified from pFA-GFP-URA3 (35) with primers GTW11 and GTW12. The resulting PCR product was cut with NsiI and EcoRV and ligated into the same sites of pUC-attR2. The resulting plasmid was then cut with SalI and NsiI and the GFP-bearing fragment ligated into the same sites of pCA-Dest1100, yielding pCA-Dest1102. For cloning of the TAP-tag, the TAP-tag coding region was amplified from CIp10-P*_PCK1_*-GTW-TAPtag (11) with primers GTW15 and GTW16. The PCR product was cut with BsrGI and NsiI and cloned in the same sites of pUC-attR2. The resulting plasmid was digested with SalI and NsiI, and the TAP-containing fragment ligated in the same sites of pCA-Dest1100, yielding pCA-Dest1103.

#### Promoters

We replaced P*_TET_* with either P*_PCK1_* (obtained from CIp10-P*_PCK1_*-GTW-TAPtag, (11)), yielding the pCA-Dest12xx series; P*_TDH3_* (cut from pKS-P*_TDH3_*; P*_TDH3_* was PCR amplified from SC5314 genomic DNA with oligos SZ11 and SZ12, and the fragment inserted into pBluescript-KS (+) cut with XhoI and EcoRV) yielding the pCA-Dest13xx series; or P*_ACT1_*, cut from CIp10::P*_ACT1_*-gLUC59 (37), yielding the pCA-Dest14xx series.

#### Selection markers

In all plasmids except the P_*TET*_ series, the auxotrophic marker *URA3* was replaced with the *NAT1* marker conferring nourseothricin resistance. Briefly, a XbaI-DraIII fragment or SpeI-NaeI fragment was cut from pUC-NAT1 and inserted into the similarly cut vectors. pUC-NAT1 was built by amplifying *NAT1* under the control of P*_TEF1_* promoter with oligonucleotides UZ50 and UZ51, and using pFA6-SAT1 (38) as a template, and cloning the AatII-cut fragment into the same site of pUC18.

The 49 expression vectors are available from Addgene (https://www.addgene.org).

### Validation of 15 expression plasmids

#### Construction of C. albicans UME6-overexpression strains

Detailed methods for the transfer of *C. albicans* ORFs from pDONR207 into the expression plasmids as well as the integration of the resulting expression plasmids at the *RPS1* locus have been described (34). Briefly, the *UME6* ORF was transferred from the entry clone into each one of 15 expression vectors by using the Gateway^™^ LR Clonase^™^ II Enzyme Mix (Invitrogen). After *E. coli* transformation, the plasmids were verified by EcoRV digest. The *URA3-* and *NAT1*-bearing expression plasmids were digested by StuI and transformed into *C. albicans* strain CEC161 or CEC369, respectively, according to Walther and Wendland (39). Transformants were selected for prototrophy or nourseothricin resistance, respectively, and verified by PCR using primer CIpUL with (i) primer CIpUR for the *URA3*-bearing plasmids and (ii) primer CgSAT1-rev for the *SAT1*-bearing plasmids, that yield 1 kb and 1.6 kb products, respectively, if integration of the OE plasmid has occurred at the *RPS1* locus.

#### Induction of the Tet-On system

Overexpression from P*_TET_* was achieved by the addition of anhydrotetracycline (ATc, 3 μg/ml; Fisher Bioblock Scientific) in YPD at 30°C (40). Overexpression experiments were carried out in the dark, as ATc is light sensitive.

#### Microscope analysis for filamentation

Cells were observed with a Leica DM RXA microscope (Leica Microsystems) with an x40 oil-immersion objective.

### Analysis of TAP- and 3HA-tagged proteins by Western blots

For the P*_TET_*, P*_TDH3_* and P*_ACT1_* strains, a 30 ml culture in YPD or YPD+ATc3 was inoculated at OD600 = 0.2 with a freshly grown colony and incubated at 30°C with shaking until OD600 reaches ∼1. For the P*_PCK1_* strain, cells were scraped off the agar of Petri dish cultures and resuspended in 1 ml dH_2_O after centrifugation. The cell suspension was used to inoculate a 40 ml SD culture at OD600 = 0.2.

20 ODs of exponentially growing cells were collected by centrifugation and resuspended in lysis buffer (9M Urea, 1.5% w/v dithiothreitol, 2-4% w/v CHAPS, and 1.5 M Tris pH9.5). Homogenization of the cells was achieved using a FastPrep bead beater. After homogenization, the lysed cells were centrifuged at 13,000 rpm for 10 min at room temperature, and the supernatant containing the solubilized proteins was used directly or stored at −80 °C.

Proteins were separated on an Invitrogen 8% NuPage gel, transferred onto nitrocellulose. TAP- and 3HA-tagged proteins were detected using peroxidase-coupled anti-peroxidase and anti-HA-peroxydase antibodies (Sigma and Roche, respectively) and an ECL kit (GE Healthcare).

### Construction of opaque mating-compatible two-hybrid strains and Gateway^™^-compatible two-hybrid vectors

Mating-compatible strains were generated by deletion of MTL**a** or MTLα locus using the *SAT1* flipper cassette (38) in the two-hybrid strain background SC2H3 (32). To delete the MTL**a** locus, flanking regions were amplified with primers MTLa-5’F and MTLa-5’R, and with MTLa-3’F and MTLa-3’R. To delete the MTLα locus, homologous regions were amplified with primers MTLα-5’F and MTLα-5’R, and with MTLα-3’F and MTLα-3’R. Both fragments for each locus were cloned into pSAT1 (38), at SacI/NotI and XhoI/KpnI sites respectively. SacI/KpnI deletion constructs were transformed in SC2H3 (32) and transformants were selected on YPD medium containing 200 μg/ml of nourseothricin. Positive SC2H3a and SC2H3α strains were subsequently grown in maltose medium for induction of the recombinase and loss of the *SAT1* flipper cassette.

For induction of opaque switching, a *WOR1* fragment was amplified with primers WOR1F and WOR1R and cloned into pNIM1 vector (40) at SalI and BglII sites. SC2H3a and SC2H3α strains were transformed with pNIM1-WOR1 with selection on nourseothricin-containing medium. The resulting strains, SC2H3a-pWOR1 and SC2H3α−pWOR1, were grown in presence of doxycycline (50 μg/ml) to induce the opaque state (41). Opaque mating-compatible strains were finally transformed with prey or bait plasmid and selected on SC-ARG or SC-LEU respectively.

Bait and prey two-hybrid vectors, pC2HB and pC2HP respectively (32) were converted into Gateway^™^ destination vectors using the Gateway^™^ Vector Conversion System to generate pC2HB-GC and pC2HP-GC respectively. The genes encoding the bait protein Hst7 (Orf19.469) and the prey Cek1 (Orf19.2886) were transferred by LR reactions from the donor collection of vectors into pC2HB-GC and pC2HP-GC destination vectors respectively.

### Mating approach and protein-protein interaction detection

Opaque cells of SC2H3a and SC2H3α, expressing VP16-Cek1 (Arg+) and lexA-Hst7 (Leu+) respectively, were crossed as follows. Opaque cells of both types were mixed at 1 × 10^6^cells each in Spider medium (42) and incubated at 23°C for 24h. Resulting tetraploid cells were selected on SC-LEU-ARG medium for 48h incubation at 30°C, and transferred to SC-HIS-MET medium for protein-protein interaction detection. Chromosome loss was induced on pre-sporulation medium at 37°C for 10 days as described previously (43).

## RESULTS

### Establishment of a sequence-validated Candida albicans ORFeome

Our objective was to develop a *C. albicans* ORFeome encompassing as many of the 6,205 ORFs predicted in Assembly 21 of the *C. albicans* genome as possible (44). To this aim, forward and reverse oligonucleotides with, respectively, *attB* and *attP* sequences at their 5’ ends were synthesized for each of the 6,205 ORFs (**Supplemental Table S1**), used in independent PCR reactions with genomic DNA of *C. albicans* strain SC5314 (45) and the resulting PCR products were cloned in pDONR207 using Gateway^™^ recombinational cloning (30). The resulting plasmids were then subjected to Sanger sequencing at the 5’ and 3’ ends of the cloned ORFs and to Illumina sequencing throughout the cloned ORFs.

Among the 6,205 ORF sequences (from Assembly 21) used for oligonucleotide design, 6 are no longer present in Assembly 22 while 20 are annotated as mitochondrial genes in Assembly 22. Among the 6,179 remaining ORFs, 5,102 (82.6%) were successfully cloned and full-length sequence-validated. Sequences were validated by comparing the entire sequence of each cloned ORF against the reference sequences for haplotype A and B of the *C. albicans* SC5314 genome available at the Candida Genome Database (CGD; version_A22-s07-m01-r18; (46)). We defined 4 groups: (i) 4,209 sequences that are 100% identical with a reference sequence (**No SNP, no DIP**; where SNP stands for Single Nucleotide Polymorphism and DIP stands for Deletion/Insertion Polymorphism); (ii) 116 sequences that do not carry any SNP but carry DIPs (**No SNP, DIPs**); (iii) 682 sequences that carry SNPs but no DIP (**SNPs, no DIP**); and (iv) 95 sequences that carry both SNPs and DIPs (**SNPs and DIPs**) (Fig. 1A; **Supplemental table S1)**.

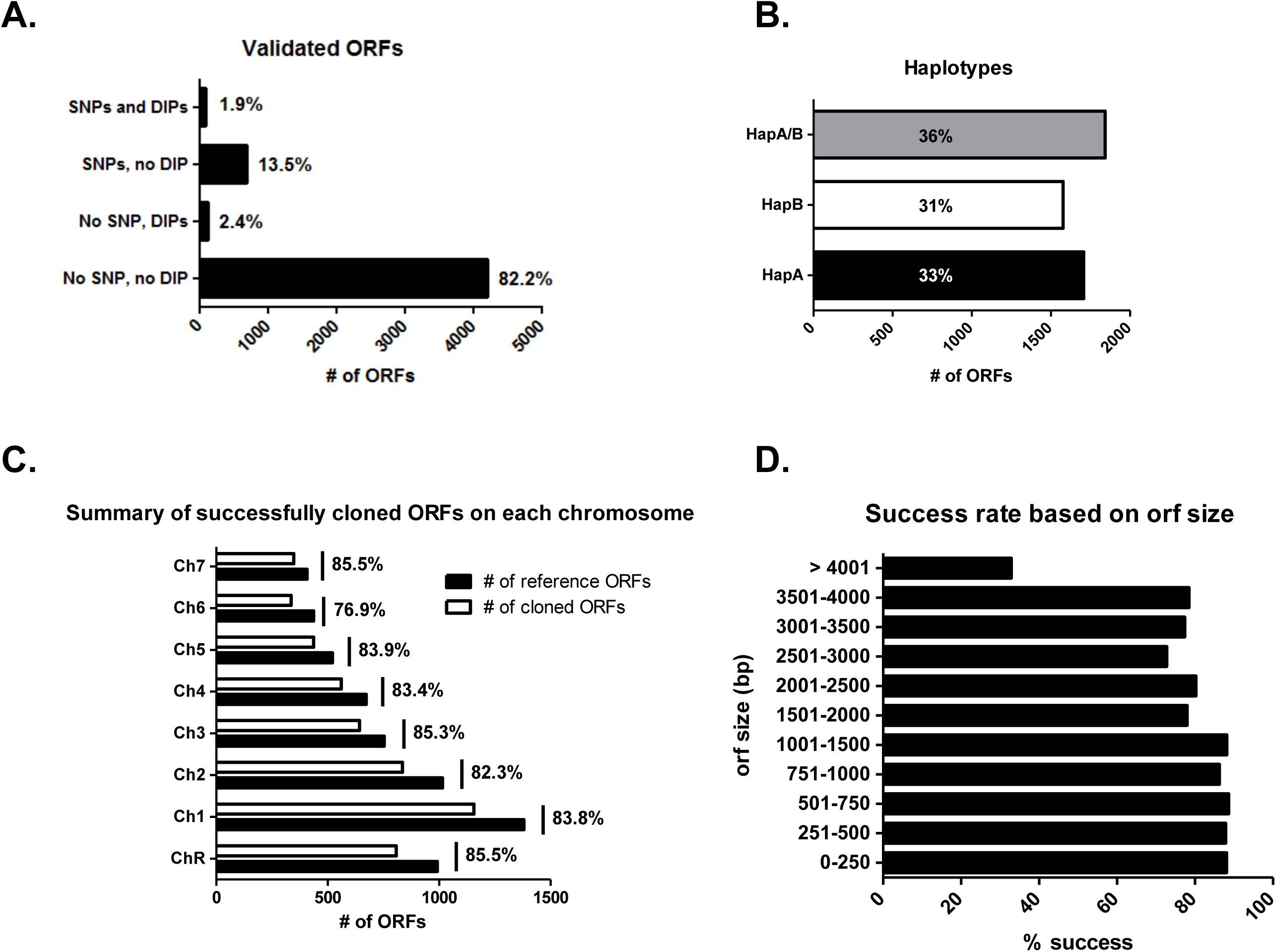

Overall, 4,209/6,179 variation-free *C. albicans* ORFs (68.1%) are now available to the community. For 35.9% of the validated ORFs, there was no difference between the two alleles in the reference sequences (**HapA/B;** Fig. 1B). When nucleotide differences were present in the two alleles of the reference ORF sequence, we did not notice any bias in the haplotype that was cloned: haplotype A ORFs were cloned in 33% of cases (**HapA;** Fig. 1B) and haplotype B ORFs were cloned in 31% of cases (**HapB;** Fig. 1B). On chromosomes R, 1, 2, 3, 4, 5 and 7, ORF cloning was uniformly successful, with a cloning success rate ranging from ∼82% to ∼86%. Chromosome 6 ORFs were slightly underrepresented (76.9%) (Fig. 1C). Although ORFs up to 1.5kb could be cloned with a success rate around 87% and ORFs between 1.5 and 4kb could be cloned with a success rate around 77%, we observed a decreased success rate for the longer ORFs (Fig. 1D).

The analysis of the 6,880,620 nucleotides of the 5,102 sequence-validated ORFs revealed 2077 SNPs. These nucleotide substitutions could be real SNPs between our strain and the reference strain, or due to poor quality of some reference sequences, or mutations in the primers, or misincorporation by the DNA polymerase. Overall, the resulting maximum error rate amounts to 3 × 10^−4^or 1 SNP every 3,312 cloned nucleotides. Notably, the *C. albicans* SC5314 genome sequence available at CGD (version_A22-s07-m01-r18) contained 272 ORFs with sequence ambiguities for at least one of the alleles. These ambiguities included stretches of ‘Ns’ or IUPAC code nucleotides (Y, R, S, M, K, W). The ORFeome-generated sequencing data resolved sequence ambiguities for 110 of these ORFs (**Supplemental Tables 3 and 4**). In this study, we also detected recombinant haplotypes for 116 of the cloned ORFs (**Supplemental Table 5**) that display characteristics of both reference haplotypes.

Taken together, our study provides an almost comprehensive, extremely high quality *C. albicans* ORFeome.

### CandidaOrfDB: a database for the C. albicans ORFeome

CandidaOrfDB was created to integrate the data of the *C. albicans* ORFeome project and is available at http://candidaorfeome.eu. CandidaOrfDB enables the scientific community to search for availability and quality of the clones. All data pertinent to the cloning process are stored in the database. This information includes the name and sequence of the primers used to amplify the ORFs from *C. albicans* SC5314 genomic DNA, the coordinates of the primers in their 96-well storage plates, and any sequencing information available on the clones. SNPs and DIPs, their location in the ORF and their influence on the corresponding amino acid sequences are shown (Fig. 2). Nucleotide sequence of the cloned ORF and the corresponding amino acid sequence are displayed with highlights of SNPs or DIPs relative to the closest haplotype sequence (Fig. 2). The reference sequence data have been extracted from CGD (46). The database can be queried through the ORF number (orf19.xxxx or 19.xxxx; (5)), the Assembly 22 name (Ci_XXXXXW or Ci_XXXXXC; (5)) or the gene name.

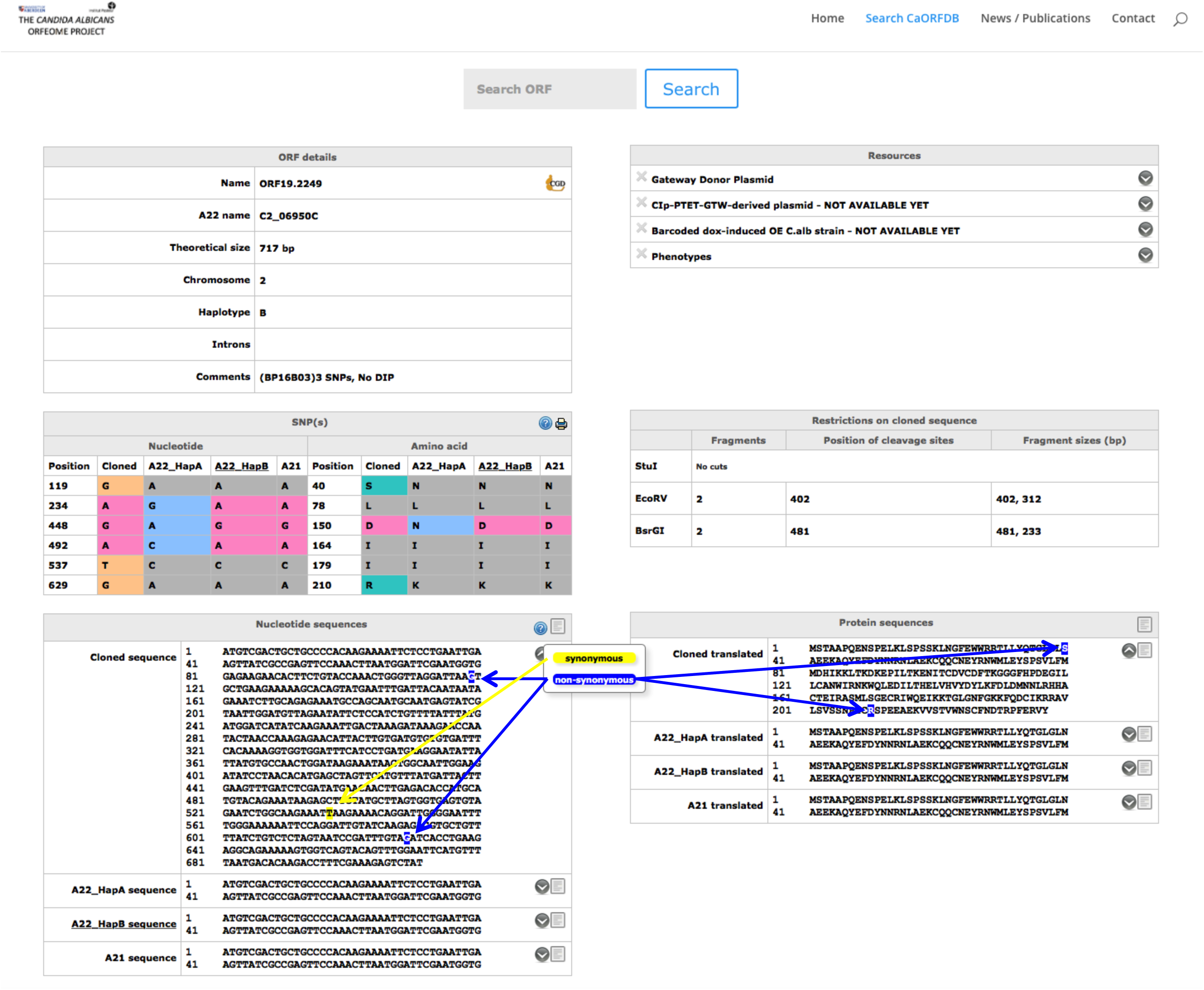

### A collection of destination vectors for exploiting the C. albicans ORFeome

The *C. albicans* ORFeome described above was developed in pDONR207 as this allows subsequent transfer of the cloned ORFs in Gateway^™^-adapted destination vectors. While destination vectors for ORF expression in hosts such as *E. coli* or *S. cerevisiae* are available (47,48), only a few destination vectors for ORF expression in *C. albicans* have been reported (11). Therefore, we set out to establish a collection of Gateway^™^-adapted destination vectors for constitutive or conditional expression of untagged or tagged ORFs in *C. albicans*. Our primary collection of 15 *C. albicans* destination vectors derived from CIp10S (11) provides a choice of two promoters, either the constitutive promoter P*_TDH3_* or the inducible promoter P*_TET_*, and the option for N- or C-terminal fusion to various epitope tags (3xHA or TAP) (**Table 2** and Fig. 3). Integration of the destination vectors and their derivatives at the *RPS1* locus in the *C. albicans* genome is promoted by StuI or I-SceI linearization. These CIp10-derived vectors carry either the auxotrophic marker *URA3* or the *NAT1* marker that confers resistance to nourseothricin and can be used with clinical isolates. However, P*_TET_*-bearing plasmids cannot carry the *NAT1* marker since the transactivator needed for tetracycline-mediated induction of P*_TET_* is borne on the *NAT1*-carrying pNIMX plasmid (11). All vectors harbor a so-called spacer sequence (SP) originating from the *E. coli* kanamycin resistance gene, for barcode insertion and subsequent Illumina-based barcode sequencing. A second set of 34 plasmids was also generated with the constitutive promoters P*_ACT1_* or the inducible promoter P*_PCK1_*, and the option for N- or C-terminal fusion to GFP (**Table 2**). All 49 destination vectors were designated with a standardized nomenclature, pCA-DESTijkl, whereby i stands for the transformation marker (1, *URA3*; 2, *NAT1*); j stands for the promoter (1, P*_TET_*; 2, P*_PCK1_*; 3, P*_TDH3_*; 4, P*_ACT1_*); k stands for N-terminal tagging (0, no tag; 1, 3xHA; 2, GFP; 3, TAP); and l stands for C-terminal tagging (0, no tag; 1, 3xHA; 2, GFP; 3, TAP) (**Table 2** and Fig. 3). All destination vectors are available from Addgene (https://www.addgene.org).

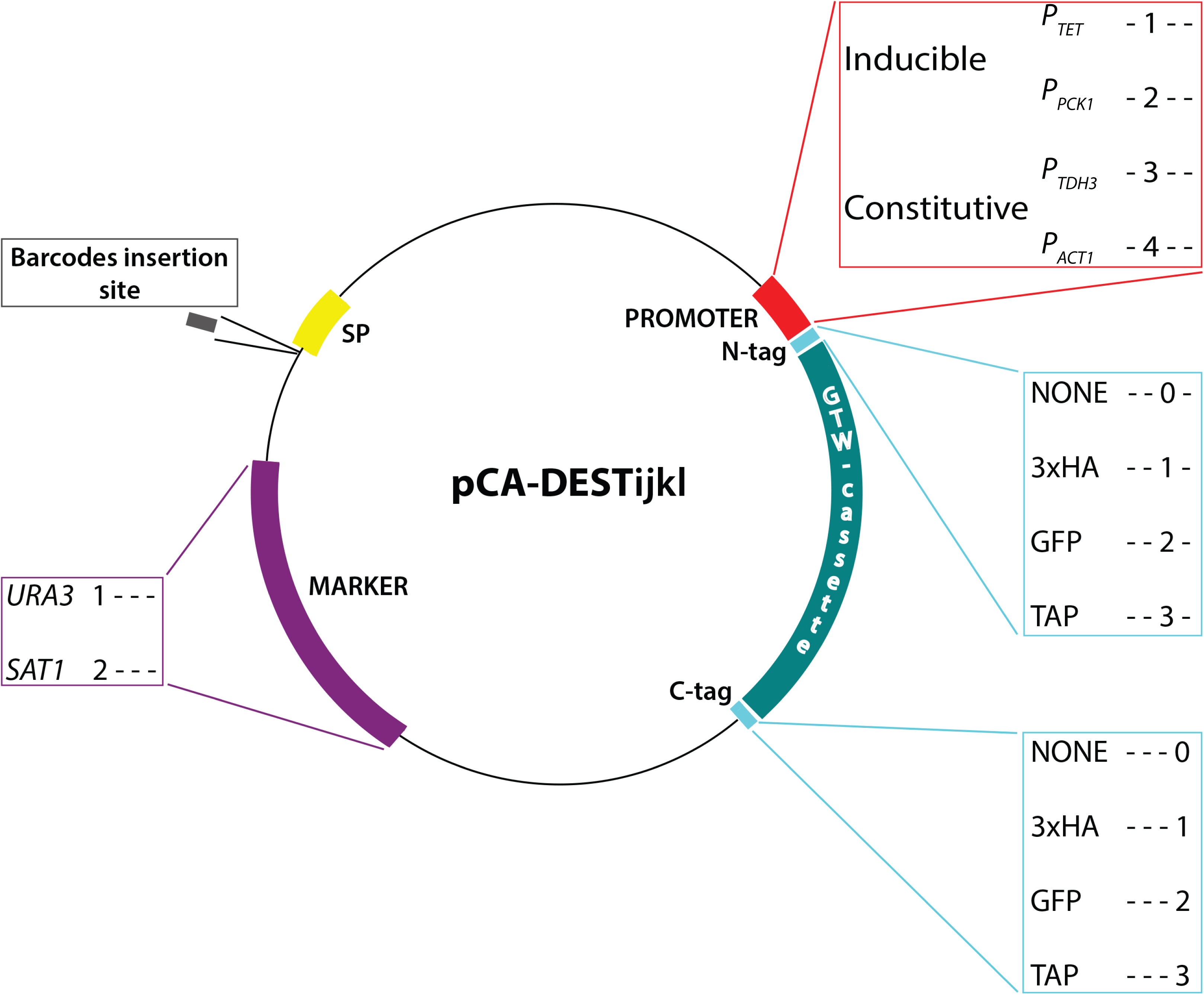

In order to validate that the primary set of 15 destination vectors could drive constitutive or conditional expression of untagged or tagged ORFs, the *UME6* ORF was transferred into each of them using LR Clonase^™^. *UME6* encodes a transcription factor whose overexpression has been shown to force filamentation in *C. albicans* (49,50). The validation criteria for the destination vectors were (i) success of the LR reaction, (ii) integration in the *C. albicans* genome at the *RPS1* locus, (iii) filamentation phenotype and/or (iv) detection of the 3xHA- or TAP-tagged Ume6 protein by western blot. The LR reaction was successfully used to transfer *UME6* from the pDONR207:: *UME6* plasmid to all 15 destination vectors. The resulting expression plasmids were successfully integrated at the *C. albicans RPS1* locus. The 5 strains containing P*_TET_* were induced overnight at 30°C by adding 3 μg/ml anhydrotetracycline (ATc3) to the culture medium, while the 10 strains containing P*_TDH3_* were grown overnight at 30°C in YPD. As described in Chauvel *et al*. (11), overexpression of Ume6 led to hyperfilamentation. This phenotype was observed for all strains, regardless the tags (Fig. 4A and 4B). 3xHA- or TAP-tagged Ume6 was detected in the relevant strains and under the appropriate growth conditions (Fig. 5A and 5B).

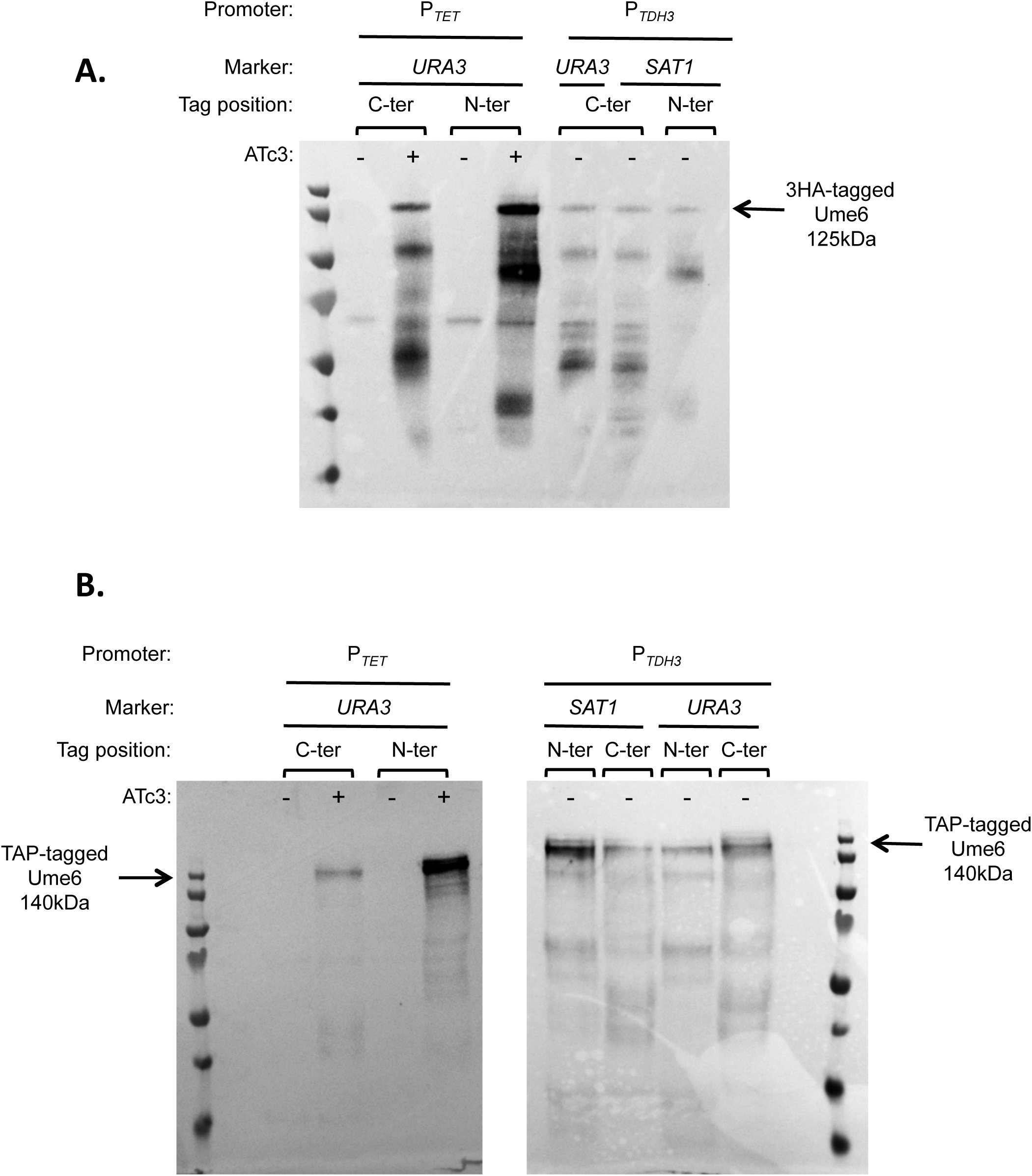

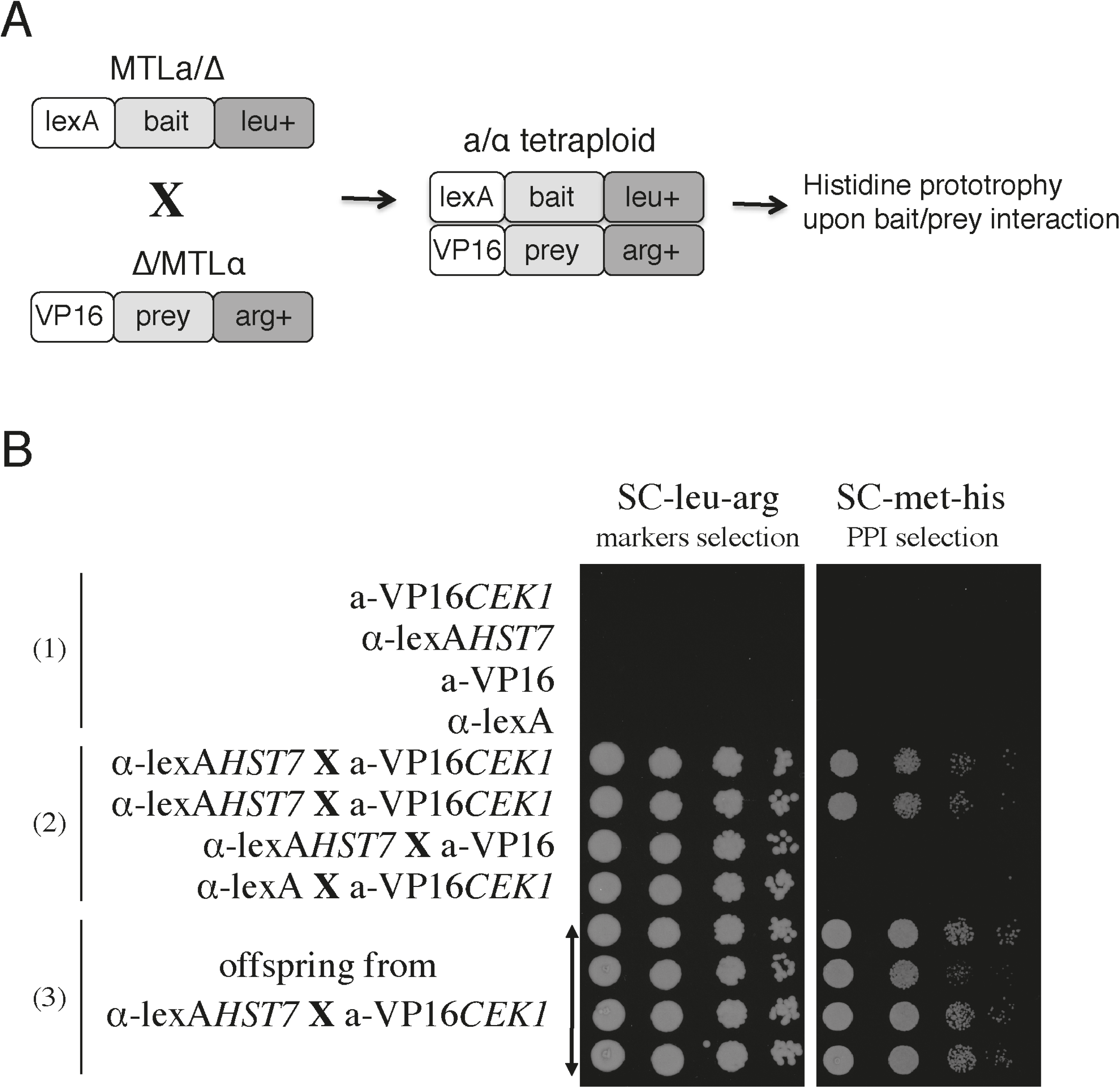

Taken together, these data present a collection of 49 available destination vectors and validate 15 of those for application with the *C. albicans* ORFeome.

### Proof of concept of a two-hybrid matrix approach of protein-protein interaction detection via mating in C. albicans

The availability of the ORFeome collection is a prerequisite to large, systematic two-hybrid (2H) screenings in *C. albicans*. The Candida 2H (C2H) system was developed for targeted one to one protein-protein interaction detection (32). Hence, there was a need to engineer the system for high throughput screening based on rapid and convenient recombination cloning systems for the generation of prey/bait arrays of proteins, and to use a mating approach to circumvent the low efficiency of transformation of this fungus. The former was obtained by adapting the prey and bait recipient vectors for Gateway^™^-mediated transfer of the ORFeome library, yielding pC2HB-GC and pC2HP-GC, respectively. The latter involved the combined processes of white-opaque switching and the preparation of mating-compatible 2H strains. The pleiomorphic fungus *C. albicans* is characterized by a parasexual cycle, with the possible mating of diploids, and generation of tetraploids, the absence of meiosis and the reversion to the diploid state by chromosome loss (reviewed in (51)). An essential pre-requisite to mating in *C. albicans* is the phenotypic switch from white to opaque, a stable but reversible switch regulated by environmental stimuli such as low temperatures, carbon dioxide and nutrients. The epigenetic switch is tightly linked to mating since the a1/α2 complex, encoded at the mating type like (*MTL*) loci, acts as a repressor of opaque switching (52). In that context, the two-hybrid strain SC2H3 originating from the diploid a/α SN152 strain was deleted for the *MTL*a or *MTLα* locus to construct SC2H3α and SC2H3a strains. The efficient switch to opaque in the hemizygote strains was achieved by the doxycycline-controlled expression of *WOR1*, a major regulator of the white to opaque switch (41,53).

A proof of principle of the whole procedure for protein-protein detection is shown in Figure 6. The MAP kinase kinase Hst7 was used as bait and linked to the DNA binding domain lexA, as part of the pC2HB-GC vector. Similarly, the Cek1 kinase was expressed as a prey protein, fused to the activation domain VP16, component of the pC2HP-GC vector. pC2HB-Hst7 and pC2HP-Cek1 were transformed into opaque SC2H3α and SC2H3a strains, respectively. Mating of the resulting transformants was achieved through selection for leucine and arginine prototrophy. Upon interaction of the two proteins, the reconstituted transcriptional module promoted the expression of the *HIS4* marker, allowing only the mating-derived strains to grow on a histidine-free medium, whereas the original diploid strains could not (Fig. 6B). Through a mechanism of chromosome loss, the crossed tetraploid strains return back to a diploid state. We showed that the plasmids were stably expressed in the diploid offspring strains as these strains conserved their ability to grow in absence of arginine and leucine, indicating the presence of the prey and bait vectors respectively. In addition, all these strains were able to grow in absence of histidine, indicative of the targeted protein-protein interaction.

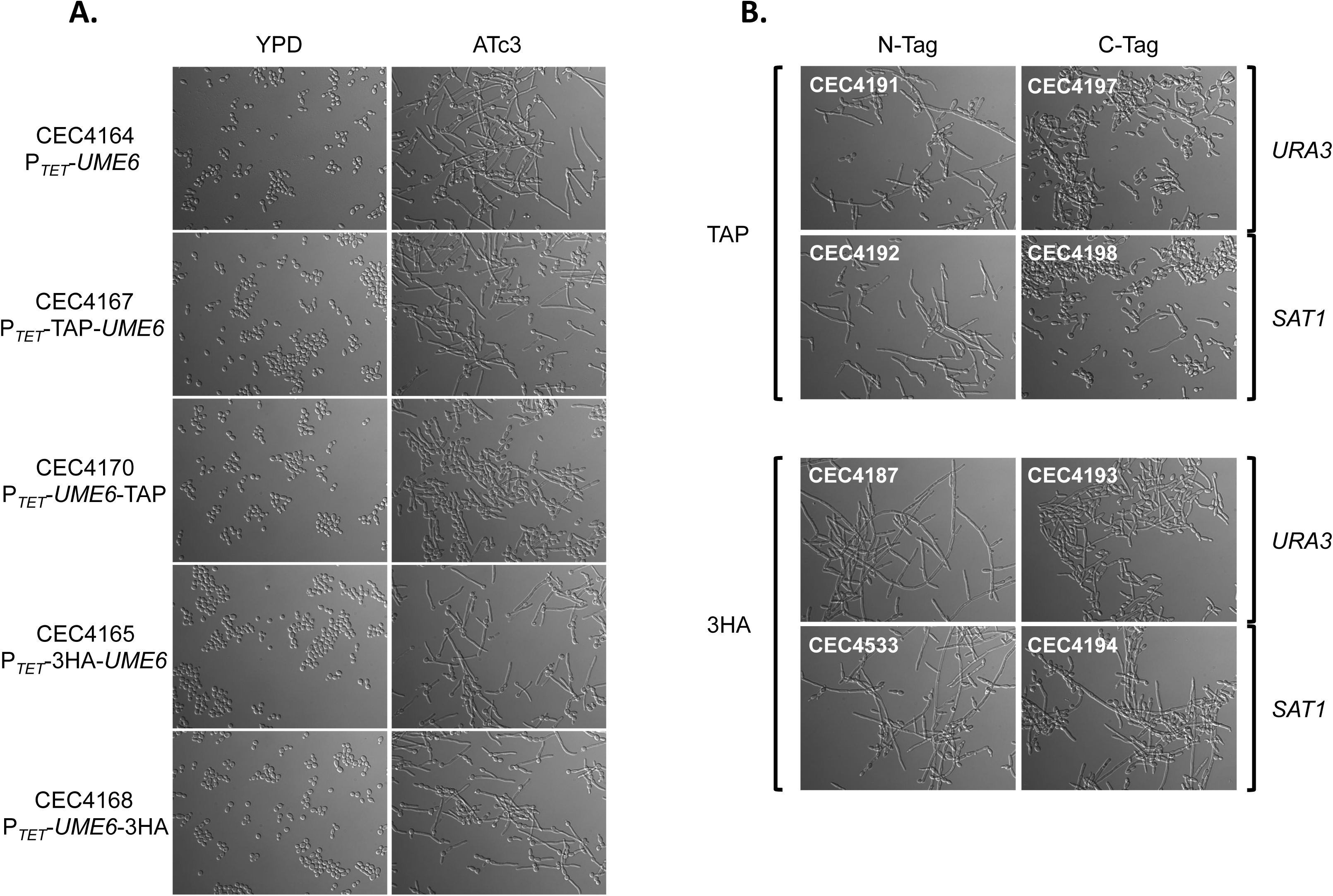

## DISCUSSION

A partial *C. albicans* Gateway^™^-adapted ORFeome (ORFeomeV1; 644 ORFs) and the corresponding *C. albicans* overexpression strains have already been successfully used to investigate morphogenesis, biofilm formation and genome dynamics in *C. albicans* (11,31,54). In this study, we have now generated the *C. albicans* ORFeomeV2 within the Gateway^™^ recombination cloning system, as well as a collection of destination vectors suitable for expression in *C. albicans*.

This new *C. albicans* ORFeome resource encompasses 5,102 ORFs (83% of the annotated ORFs) and will be made available to the community upon request. The resource is supported by the CandidaOrfDB database, providing information on the individual plasmids and their sequences. The *C. albicans* ORFeomeV2 is of unprecedented quality, as all ORFs in the collection are fully sequenced, which remains an exception in large-scale ORFeome projects where ORFs are rarely sequenced in their entirety and only the 5’- and 3’-ends of the cloned ORFs are sequenced to confirm identity and the absence of frameshift mutations in the primers (15,16,18,21,23,29). Given these high standards, the success rate of 83% that we achieved with the *C. albicans* ORFeomeV2 is quite remarkable.

In eukaryotes, ORFeomes are usually cloned from full-length cDNA libraries to account for the presence of introns and therefore splicing variants. However, only 6% of *C. albicans* genes have introns (55), rendering the generation of the *C. albicans* ORFeome more straightforward than for pluricellular eukaryotes. Indeed, ORFs were simply amplified from the genomic DNA of the reference strain SC5314 and therefore, 360 ORFs in the *C. albicans* ORFeomeV2 harbor introns, a matter that should be taken into consideration when expressing ORFs in a prokaryotic host and, possibly, other eukaryotic hosts than *C. albicans*. A second aspect to be taken into consideration when using the ORFeome in hosts other than *C. albicans* is the unusual codon usage in this species (56). Indeed, in *C. albicans*, CUG is decoded as a serine instead of a leucine the majority of the time. Hence, improper translation in hosts with a standard genetic code may distort protein structure and function.

Sanger sequencing of both 5’ and 3’ ends was performed to confirm ORF identity and exclude clones containing primer or recombination errors. Previous studies have reported mutation rates in primers of 3-10% (26,29). Similar rates were observed in this work. Unlike other Gateway^™^ recombinational cloning projects (15,29), we did not see major differences in cloning efficiency for ORFs up to 4 kb. A decrease in success rate was observed only for ORFs >4 kb. This size bias could be attributed to an increased difficulty to amplify the ORF by PCR, a reduced efficiency of the Gateway^™^ BP Clonase^™^ reactions with long PCR products, and the error rate of the PCR polymerase, which increases with longer products. In addition, we also corrected the sequences of 110 ORFs with sequence ambiguities (N-tracts and IUPAC code nucleotides) according to the Candida Genome Database and identified 116 ORFs that display a recombinant haplotype. Recombinant haplotypes could be explained by template switching during PCR cycles or by incorrectly phased SNPs in reference sequences. This could be addressed by performing long-read sequencing of the *C. albicans* SC5314 genome.

However, many challenges remain to be addressed. For heterozygous genes, only one allele is present in the ORFeome. Because functional differences have been reported between the 2 alleles of a heterozygous gene (57,58), it would be relevant to also clone and validate the second allele in these instances. In addition, oligonucleotides have been designed based on annotation of the haploid set of Assembly 19 of the *C. albicans* genome. As a consequence, genes of the *MTLα* locus have not been included in the ORFeome. Despite our efforts, the collection is still missing 1,081 clones. The next version of the *C. albicans* ORFeome could extend gene coverage by adding the missing ORFs, allelic variants or strain-specific variations.

ORFeomes are essential to bridge the gap between genome annotation and systems biology and allow large-scale gene and protein characterization. The development of a collection of *C. albicans* constitutive or conditional overexpression strains is one of the applications of the *C. albicans* ORFeome that we have successfully explored (11,31,54). In this respect, the *C. albicans* ORFeome project is currently completing its second and third phases, which consist of transferring the 5,102 cloned ORFs into barcoded destination vectors and generating a collection of *C. albicans* barcoded overexpression mutants, each mutant carrying one of the 5,102 cloned ORFs under the control of the inducible P*_TET_* promoter. CandidaOrfDB already provides information on the available overexpression plasmids and strains. In this study, we have paved the way for a second application of the *C. albicans* ORFeome through the development of tools for a 2H matrix approach of protein-protein interaction detection via mating in *C. albicans*. Our data show that mating of diploid *C. albicans* expressing a bait fused to the LexA DNA binding domain and a prey fused to the VP16 activation domain, respectively, allows protein-protein interactions to be tested. In *S. cerevisiae*, large-scale 2H screens can be performed in a matrix design whereby haploid strains expressing baits and preys are mated and protein-protein interactions are scored in the resulting diploids. Our toolkit now enables the implementation of such a matrix design for large-scale 2H screens in *C. albicans*. To this aim, a collection of 1500 prey clones has already been generated. As mentioned above, *C. albicans* codon usage is unusual, which limits the development of applications of the *C. albicans* ORFeome in species that use standard decoding such as *S. cerevisiae*. Our development of tools for large-scale 2H screens in *C. albicans* circumvents this limitation, opening the path for the characterization of the *C. albicans* interactome. Defining the *C. albicans* interactome will undoubtedly impact our understanding of *C. albicans* pathogenesis in humans.

In summary, high-level gene coverage coupled with the versatility of the Gateway^™^ recombinational cloning, *C. albicans-* adapted functional genomics tools and full access to clones, make the *C. albicans* ORFeomeV2 a unique and valuable resource for the scientific community that should greatly facilitate future functional studies in *C. albicans*.

## ACKNOWLEDGEMENT

We are grateful to other members of our groups for their support throughout the development of the *C. albicans* ORFeome.

## FUNDING

This work has been supported by a Biomedical Resources grant from the Wellcome Trust (The *Candida albicans* ORFeome project, 088858/Z/09/Z to CAM and CD), the European Commission (FINSysB, PITN-GA-2008-214004 to CD), the Agence Nationale de la Recherche (KANJI, ANR-08-MIE-033-01 to CD; CANDICOL, ANR-10-01 to CD) and the Interuniversity Attraction Poles Programme initiated by the Belgian Science Policy Office for funding (IAP P7/28 to PVD). CAM would like to acknowledge support from a Medical Research Council New Investigator Award (G0400284) and the MRC Centre for Medical Mycology and the University of Aberdeen (MR/M026663/1). VC was the recipient of a PhD fellowship in the framework of the FINSysB consortium (PITN-GA-2008-214004). AN was the recipient of a PhD fellowship from the DIM-MalInf Région Ile-de-France. EP was the recipient of a post-doctoral fellowship in the framework of the *C. albicans* ORFeome project (WT088858MA). TR was the recipient of a postdoctoral fellowship in the framework of the NPARI consortium (LSHE-CT-2006-037692). UZ was the recipient of a postdoctoral fellowship in the framework of Programme Fungi from Institut Carnot-Pasteur Maladies Infectieuses. SZ was the recipient from post-doctoral fellowships in the framework of the FINSysB (PITN-GA-2008-214004) and KANJI (ANR-08-MIE-033-01) consortia. HT was the recipient of a KU Leuven CREA grant (2011-2013). High throughput sequencing has been performed on the Genomics Platform, member of France Génomique consortium (ANR10-INBS-09-08). The funders had no role in study design, data collection and analysis, decision to publish, or preparation of the manuscript. We acknowledge support from the French Government’s Investissement d’Avenir program (Laboratoire d’Excellence Integrative Biology of Emerging Infectious Diseases, ANR-10-LABX-62-IBEID).

